# Decoupling cell size homeostasis in diatoms from the geometrical constraints of the silica cell-wall

**DOI:** 10.1101/2023.02.08.527611

**Authors:** Diede de Haan, Nahuel-Hernan Ramos, Assaf Gal

## Abstract

- Unicellular organisms are known to exert tight control over their cell size. In the case of diatoms, abundant eukaryotic microalgae, the layout of the rigid silica cell wall imposes geometrical restrictions on cell size. A generally accepted theory states that the need to fit any new silica element into a previously formed structure causes a reduction in size with each vegetative division cycle, until cell size restoration is achieved by a switch to another life-cycle stage. Nevertheless, several reported exceptions cast doubt on the generality of this theory.
- Here, we monitored clonal cultures of the diatom *Stephanopyxis turris* for up to two years, recording the sizes of thousands of cells, in order to follow the distribution of cell sizes in the population.
- Our results show that all *S. turris* cultures above a certain size threshold undergo a gradual size reduction, in accordance with the postulated geometrical driving force. However, once the cell size reaches a lower threshold, a constant size range is maintained by different cellular strategies.
- These observations suggest two distinct mechanisms to regulate the cell size of diatoms, reduction and homeostasis. The interplay between these mechanisms can explain the behavior of different diatoms species in various environments.

## Introduction

Diatoms are a group of eukaryotic microalgae, highly diverse in terms of morphology, cell size, and aquatic habitat (Falciatore and Mock, 2022). A unifying characteristic of diatoms is their ability to precipitate silica to form intricately shaped cell walls, called frustules (Kroger and Poulsen, 2008; Hildebrand, Lerch and Shrestha, 2018). The general layout of the frustule is conserved among diatom species, and consists of two halves. Each half is composed of a valve and a series of girdle bands. The valves are richly ornamented structures, located at opposite sides of the cell, and have species-specific morphologies with micro- and nanopatterns (Round *et al*., 2007). Appended to the rim of the valves are girdle bands, made of silica strips that encase the cell circumference. Valves and girdle bands are usually formed intracellularly, each in an individual membrane bound organelle (Reimann, 1964; Mayzel *et al*., 2021). After completion, the mature silica element is exocytosed (de Haan *et al*., 2023). Formation of new cell wall elements occurs along the cell division cycle (Fig. 1a). After vegetative cell division, each of the daughter cells inherits one-half of the parental cell wall and produces a new half in order to form a full silica covering. New valves are formed directly after cytokinesis in order to complete cell division, and girdle bands are formed later, during the G1 or G2 growth phases.

**Figure 1.**
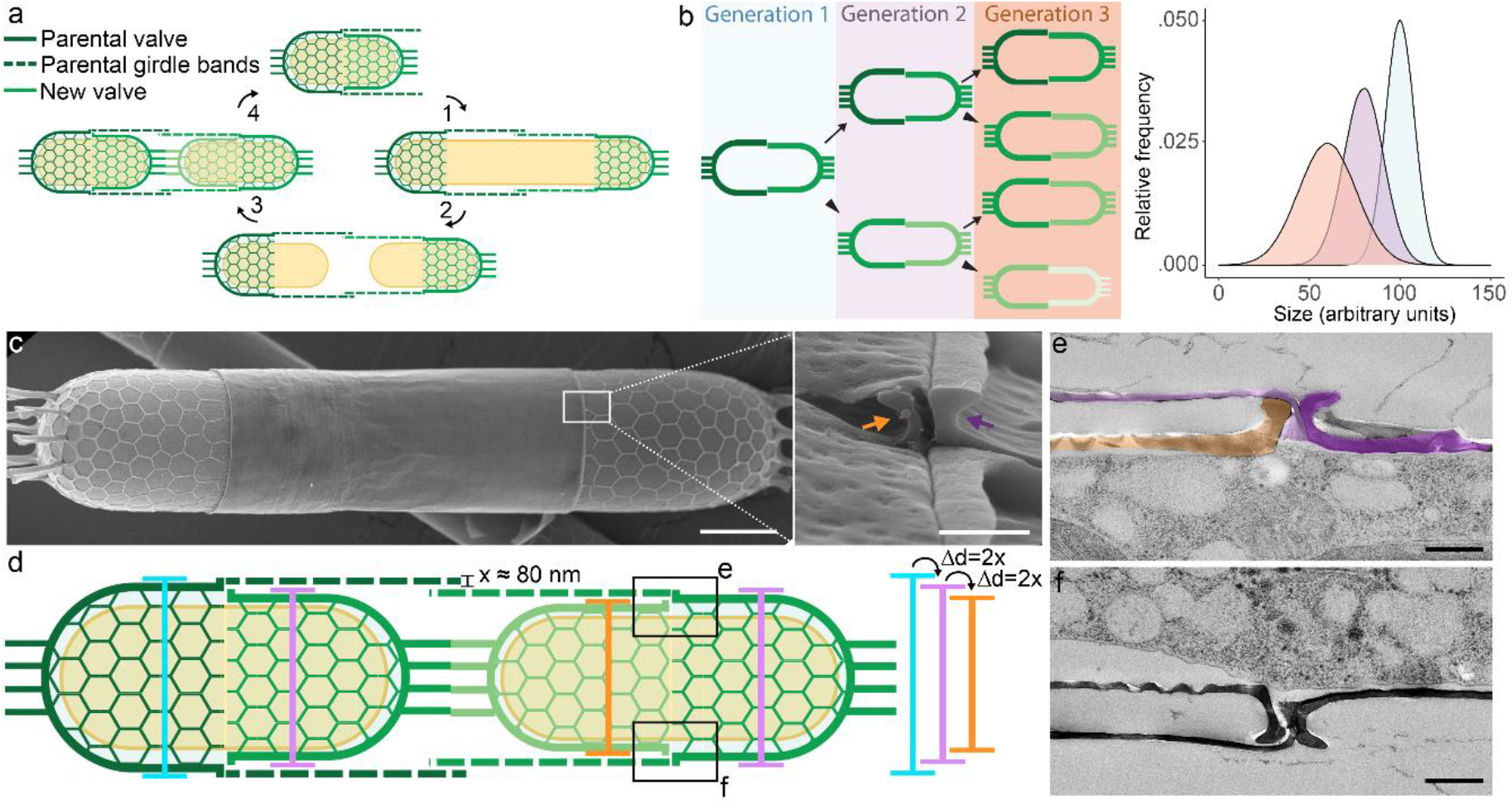
Cell wall structure and cell division in the diatom *S. turris*. (**a**) Cell cycle of *S. turris*. 1: Longitudinal growth and girdle band formation, 2: Cytokinesis, 3: Valve formation, 4: Next cell cycle. (**b**) Gradual size reduction according to the MacDonald-Pfitzer rule. One daughter cell inherits the larger half of the parental cell wall and will have the same size as the mother cell (arrows), whereas the other daughter cell inherits the smaller parental cell wall half and will be slightly smaller than the mother cell (arrowheads). As a result, the mean cell diameter of the population will decrease and the variability in diameter will increase. (**c**) SEM micrograph of two *S. turris* daughter cells, the parental valves of both cells are visible whereas the newly formed valves are covered by the girdle bands. Scale bar: 10 μm. Inset: a fractured cell shows the hook shape at valve rim. The new valve (orange arrow) is visible through a crack in the girdle bands. Scale bar: 500 nm. (**d**) Schematic of *S. turris* showing the relative diameter of parental valves and new valves. (**e-f**) TEM micrographs of the boxed areas in **d**, where parental valve (purple highlight) and new valve (orange highlight) meet under the parental girdle bands (lilac highlight). Scale bars: 500 nm

In contrast to other phytoplankton groups, diatoms do not have a single characteristic cell size but rather span many orders of magnitude across species (Litchman *et al*., 2009). Even within a single species, cells can vary greatly in their size. However, size plasticity is limited in diatoms as the silica frustules pose unique challenges for cell size control, addressed initially in the compelling MacDonald-Pfitzer hypothesis (Macdonald, 1869; Pfitzer, 1869). This hypothesis states that since each newly formed valve is formed within the bounds of the rigid parental cell wall, the new valve has to be slightly smaller than the parental valve, similar to a Petri dish (Wallich, 1860). As a result, size distributions of diatom populations should follow a pattern of decreasing mean and increasing variance over time (Fig. 1b). The escape from clonal death, due to ever-shrinking progeny, is possible through the formation of a distinct cell type, called auxospore (Kaczmarska and Ehrman, 2021). The protoplast of an auxospore is not enclosed by a rigid silica casing and thus not limited in its growth. New silica valves that are formed within the large auxospores can therefore be larger than the previous generation valves. Auxospores are usually formed through a cycle of sexual reproduction, but vegetative auxosporulation has been reported (Chepurnov *et al*., 2008).

The theory formulated by MacDonald and Pfitzer and its implications for the life history of diatoms have become a firm paradigm in the field, even though most well-known examples belong to the taxonomical group that exhibit bilateral symmetry, called pennate diatoms. A few extensively studied diatom genera, such as *Seminavis, Pseudo-nitzschia* and *Cylindrotheca* are textbook examples of the typical size reduction-restoration cycle and pheromone-induced sexuality and mating (Geitler, 1932; Chepurnov *et al*., 2002; D’Alelio *et al*., 2009; Klapper *et al*., 2021). Yet, it has been notably difficult to detect the expected size distribution cycles and life-cycle stages in both natural and laboratory populations of centric diatoms, which are radially symmetric (Table 1). Environmental factors, such as seasonal temperature and salinity fluctuations, size selective sedimentation, and parasitism have been pointed out to possibly mask underlying geometrically induced size distributions in wild diatom populations (Wimpenny, 1956; Round, 1982; Koester *et al*., 2007; Wang *et al*., 2015). In addition, numerous diatom cultures are maintained in laboratory conditions without any noticeable change of cell size, supposedly refuting the geometrical size control hypothesis. To date, these discrepancies are reconciled by stating the difficulty to observe cell size restoration in natural populations, or that some species are exceptional (Mann, 1988; Chepurnov *et al*., 2008).

**Table 1.**
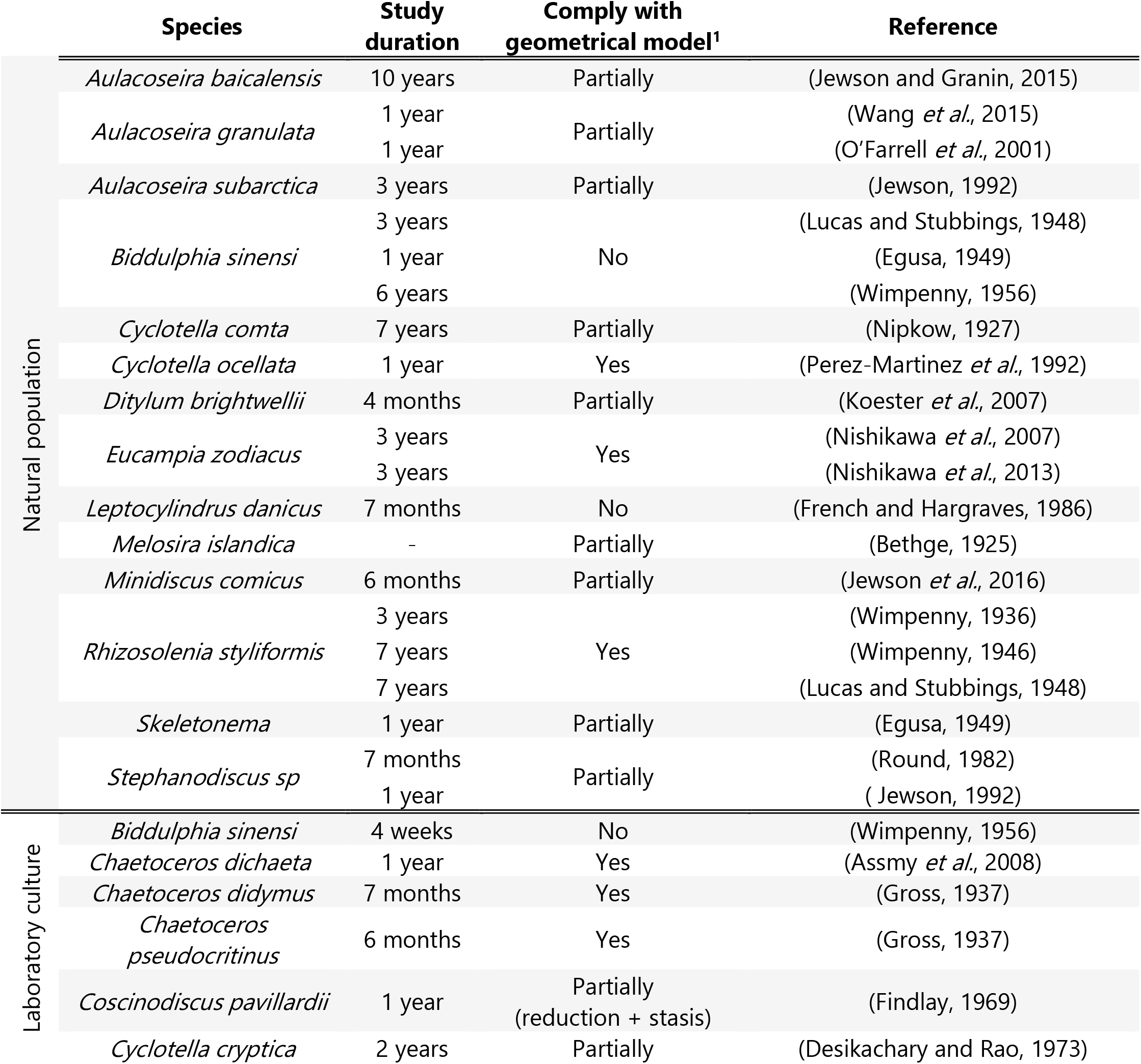
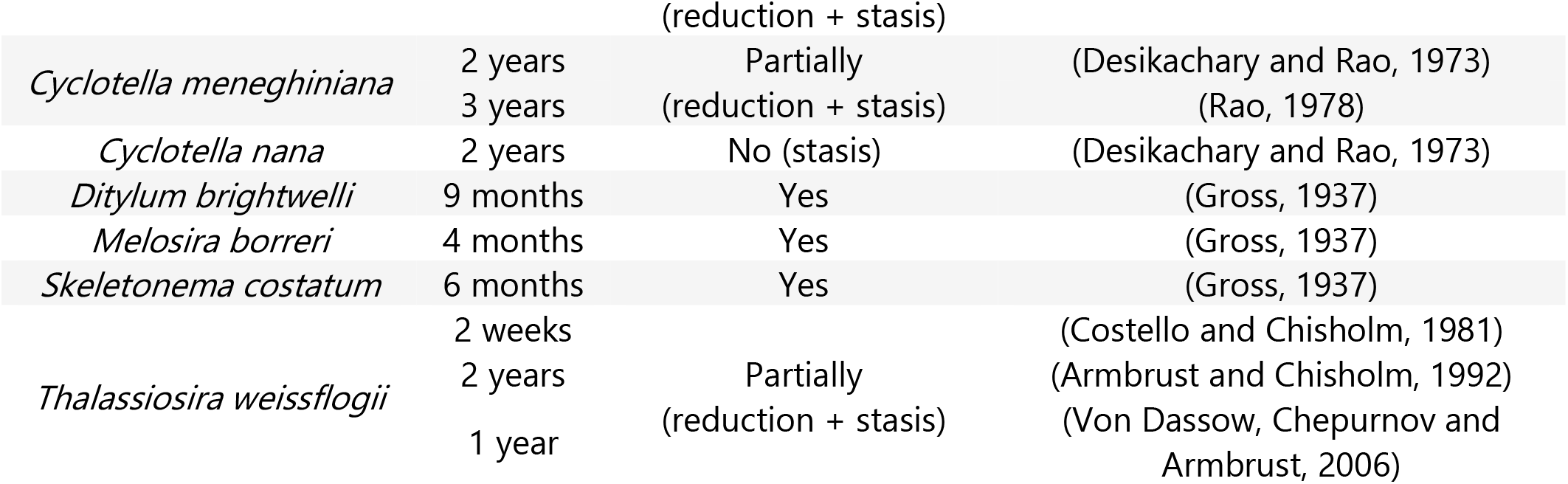
Previous studies reporting on cell size changes in centric diatoms, and their compliance with the MacDonald-Pfitzer model of cell size reduction and restoration.

Here, we design experiments that directly address the long-term progression of cell sizes in laboratory cultures of the centric diatom, *Stephanopyxis turris*. Using thousands of automated cell size measurements, we describe the process of cell size regulation over long-term culturing. Our results demonstrate that even though the predicted geometrical cell size reduction is a prevailing mechanism to reduce the size of large cells, in the long term, all cultures maintain an optimal average cell size via mechanisms that do not involve these geometrical constraints.

## Materials and methods

### Cell cultures

The cultured strain of *Stephanopyxis turris* was collected from the North Sea in 2004 and provided by the group of Prof. Eike Brunner, TU Dresden. Cell cultures were grown in sterile seawater (cFSW) from the Mediterranean, adjusted to 3.5% salinity and supplemented with f/2 nutrient recipe, under 16h/8h light/dark at 18 °C. Illumination intensity during the light period was ~90 μmoles/m^2^/s.

### Clonal isolation and maintenance

Single chains of *S. turris* cells were isolated using a 10P micropipette under a stereo microscope. After 3 transfers in drops of cFSW, single chains were placed in a 24-well tissue culture plate (Falcon, Corning, USA) filled with 1 mL cFSW and allowed to grow for 3-5 weeks, under conditions as described above. After substantial growth, the clonal cultures were used to inoculate 50 ml of fresh cFSW medium in a 250 ml Erlenmeyer flasks and again allowed to grow for a couple of weeks. Subsequently, the cultures were maintained at either a high light condition at ~90 μmoles/m^2^/s or low light condition at ~30 μmoles/m^2^/s. Grown under high light *S. turris* growth rate is about 1 division/day and cultures were maintained by weekly dilutions of 1:100. Grown under low light the growth rate is about 1 division/week and cultures were maintained by monthly dilutions of 1:10.

### Cell size measurements

For cell size measurements a 25 μl drop of cell culture was pipetted on a standard glass microscope slide and imaged using an Eclipse Ni-U upright optical microscope (Nikon, Tokyo, Japan) equipped with 10x and 20x objectives. Images for later analysis were acquired with NIS-Elements software (Nikon) using a DS-Fi3 camera (Nikon). A custom, in-house Python script was developed to facilitate high-throughput, automated cell size measurements from the acquired micrographs (Fig. S1). The script uses RGB images in .tif format as input. First, cells are segmented from the background by converting the images to grey scale and thresholding according to Otsu’s method (Otsu, 1979). Subsequently, the segmented areas in the image are filtered to remove objects that fall out of a specified size range. Finally, a range of properties of the identified cells is exported in tabulated format. For our measurements we have used mainly the minor-axis measurements of an ellipse fitted for each cell. We tested the accuracy of the automated script by comparing its output with manually measured cells sizes (Fig. S2). The script and operation instructions have been deposited on Github (https://github.com/Nahuel88Ar/Cells-Detection-).

### Simulations, data analysis and visualization

All simulations and data processing, analyses and visualizations were performed using the open source software R v 4.0.4 using Rstudio v 2022.02.3 (Team, 2013, 2020). Linear regression analysis was performed using the stats v 4.2.1 package. All data visualization was performed using the ggplot2 package and assembled in Adobe Illustrator 2021.

### Scanning electron microscopy (SEM)

Diatom cells were fixed using a solution of 2% Glutaraldehyde and 4% Paraformaldehyde in artificial seawater for 1 hour at room temperature while shaking. After 3 washes with deionized water (Milli-Q® IQ 7003 Ultrapure Lab Water System, Merck), the cells were dehydrated by washing in a graded series of ethanol. The final wash was done in 100% anhydrous ethanol overnight. The dehydrated samples were then dried in a critical point dryer using liquid CO_2_ as transitional fluid (CPD). Dried cells were placed onto a conductive carbon tape on an aluminum stub, then sputter-coated with 4 nm iridium (Safematic) and imaged on an Ultra 55 FEG scanning electron microscope (Zeiss, Germany), using 3-5 kV, aperture size 20-30 um and a working distance of about 3 mm.

### Transmission electron microscopy (TEM)

Diatoms were gently concentrated on a 5 μm filter, and washed into an Eppendorf tube into 200 μl artificial seawater. After settling, 2 μl of dense cell suspension were pipetted from the bottom of the Eppendorf tube into aluminum HPF discs (100 μm, Wohlwend GmbH, Sennwald, Switzerland) and covered with a second disc (flat). Directly after closing, the discs were loaded into a Leica ICE high pressure freezing machine and vitrified at liquid nitrogen temperature (−192 °C) at 210 MPa (Leica Microsystems GmbH, Wetzlar, Germany). Vitrified samples were kept in liquid nitrogen until freeze-substitution in a freeze-substitution device at −90 °C (EM ASF2, Leica Microsystems GmbH, Wetzlar, Germany). For 48 hours, samples were immersed in an anhydrous acetone solution containing the following combination of fixatives and stains: 0.2% uranyl acetate (UA), 0.2% osmium tetroxide (OsO4), and 2% glutaraldehyde (GA). The temperature was let to raise linearly to −20 °C over 24 hours, and then to 0 °C over 1 hour. At 0°C, samples were taken out of the FS dewar and brought to room temperature. Acetone was gradually replaced by Epon (Agar Scientific Ltd, Stansted, U.K.) using gradient concentration mixtures (10%, 20%, 30%, 40%, 60%, 80%, 100% Epon in acetone), twice a day. Epon-embedded samples were cured at 70 °C for 72 and then sectioned to 70 nm thin sections using Ultracut UCT ultra-microtome (Leica Microsystems GmbH, Wetzlar, Germany) equipped with an Ultra 45° diamond knife (Diatome Ltd, Nidau, Switzerland). Sections were mounted on copper TEM grids with a carbon support film. The sections were then post-stained with lead citrate solution. Ultrathin sections were imaged with an FEI Tecnai Spirit transmission electron microscope (FEI, Eindhoven, Netherlands) operated at 120 kV and equipped with a Gatan Oneview camera.

## Results

The life cycle of the centric diatom *S. turris* has been described in profound detail at the level of individual cells (Fig. 1a, von Stosch and Drebes, 1964; von Stosch, 1965). All aspects of the classically proposed diatom life-cycle, including vegetative cell division with size reduction, formation of gametes, auxospore formation and vegetative size restoration, have been observed in culture (von Stosch and Drebes, 1964). This makes *S. turris* an excellent candidate for a size progression study since the hypothesized constraints to which a new valve is subjected depend on species-specific geometries and the expected cell size reduction per generation can be inferred from the structural details of the cell wall geometry (Round, 1972). The silica valves of *S. turris* are ornamented with linking extensions at their apex (Fig. 1c). Through these linking extensions daughter cells remain mechanically connected after cell division, forming chains of usually 2, 4 or 8 cells. At the rim of the valves, the silica forms a slightly raised hook to which the girdle bands are appended (Fig. 1c inset, arrows). Just after its exocytosis, the new valve is positioned directly adjacent to the parental valve and completely covered under the parental girdle bands, as can be observed in stained thin sections of Epon embedded cells (Fig. 1d-f). From these arrangements, it follows that the width of the new valve is limited by the girdle bands of the parental valve. Thus, the diameter of the new valve should be smaller by at least twice the girdle band thickness (Pfitzer, 1871; Round, 1972).

To test whether the cell size of *S. turris* changes according to the MacDonald-Pfitzer hypothesis, we monitored cell size values in cultures at regular time intervals. The experimental cultures were prepared by isolating single chains with distinct cell diameters to create clonal cultures that are initially homogenous in size (Fig. 2a). Under the experimental conditions, the cells divide once per day and were diluted weekly to maintain exponential growth. Cell size was measured from a minimum of 15 microscope images, which were analyzed using in-house python scripts for automated measurements of live-cell micrographs (see Methods section and Figure S1). In brief, RGB images are converted to grey-levels and a threshold is applied to segment cells with lower pixel intensities from the background with higher pixel intensities. Then, the regions that are recognized as cells are evaluated to exclude non-ellipsoidal regions or regions that fall outside of a specified size range. Finally, morphological properties of interest (e.g. minor and major axis length) are measured for each cell in the image.

**Figure 2.**
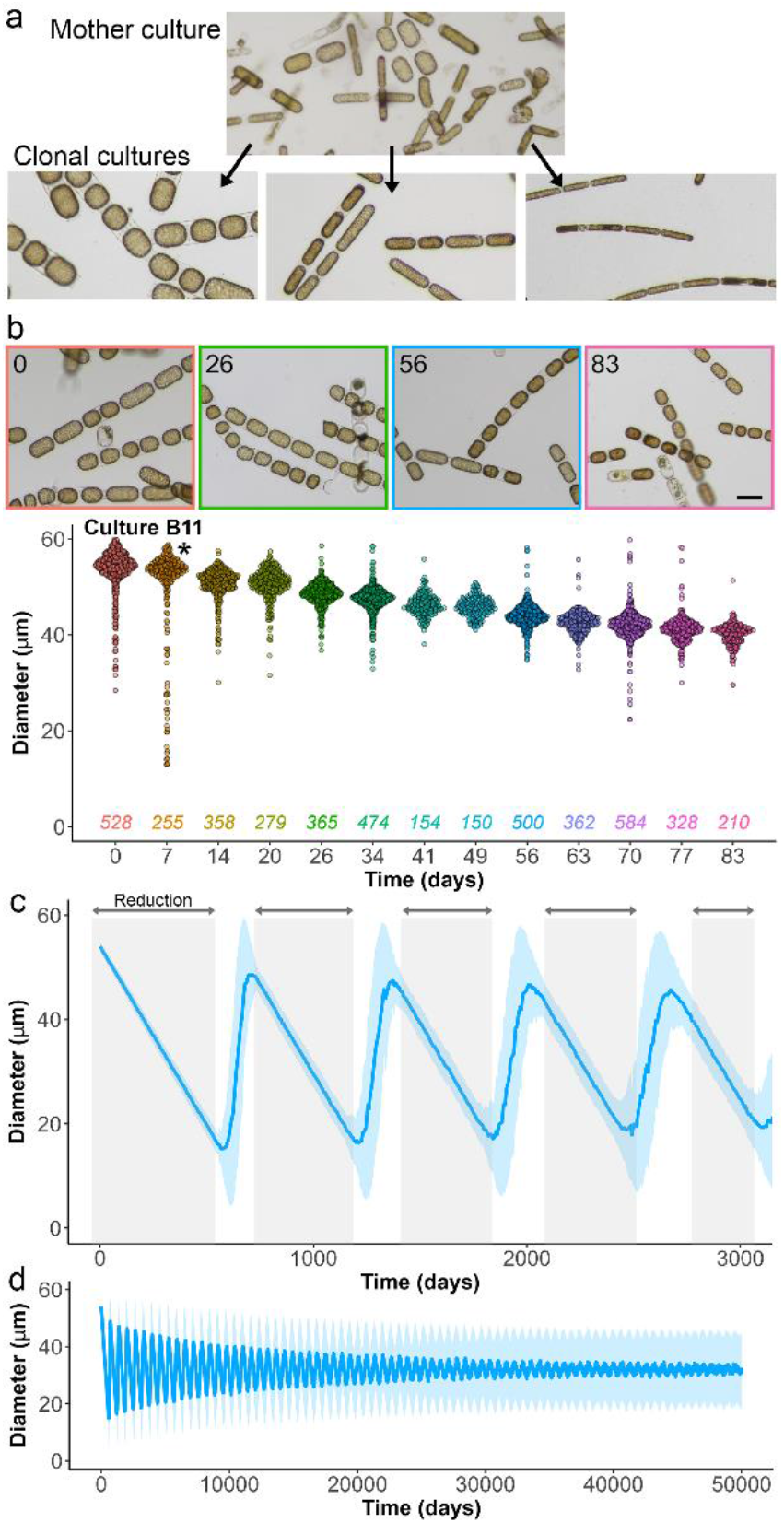
Experimental and simulated cell size change in *S. turris*. (**a**) The procedure of isolating single chains to establish clonal cultures with homogeneous cell size. Scale bar: 50 μm. (**b**) An example for the changes in cell diameter by generation, presented as ‘Bee swarm’ plots. Each point represents a single cell. Numbers below the plot indicate sample size for each time point. Micrographs above the plot show representative images of the culture at several time points during the experiment. Asterisk indicates data imaged using x20 objective, all other data points were recorded using x10 objective. (**c**) Simulated mean (line) and standard deviation (light blue area) of the diameter of *S. turris* cells for 3000 generations (at 7-generation intervals). Gray-shaded areas indicate periods of size reduction, whereas white areas indicate periods of size restoration. (**d**) Simulated culture as in (**c**) over 50000 generations

In order to evaluate the accuracy of the automated measurements we compared them with manual measurements of few data sets. This shows that the automation results give slightly higher cell diameters than the manual measurements, mostly because not all overlapping cells and impurities in the media are accurately filtered out, but correctly represent the cell size change over time (Fig. S2). Since extreme values in the datasets are the result of inaccurate filtering of non-cell objects, we chose to describe the population by its median, rather than by the mean, which gives less weight to the unrealistic outliers.

The output of this high-throughput and automated analysis, conducted on images from a typical culture that was founded by a large cell and monitored for ~3 months, shows a gradual decrease in cell size (Fig. 2b). During the first measurement, the median diameter of this culture was 54.5 μm and after 83 generations it had reduced to 40.6 μm, corresponding to an average reduction of 167 nm per generation. The culture remained unimodal throughout the experiment and the variance remained relatively unchanged over time. Note that cell volume is only slightly affected by the decrease in cell diameter, as cell length increases accordingly.

To test whether the measured size reduction in our cultures is in agreement with the MacDonald-Pfitzer hypothesis, we simulated cell size change in a highly simplified, yet relevant, computer simulation. The simulation starts with a population of 10^6^ cells that are rather homogeneous in size (for example 54±1 μm), as in the experimental setup. Each cell division produces one offspring with the same diameter as the parental cell and one offspring that is smaller than the parent by twice the girdle band thickness. *S. turris* girdle bands are around 80 nm, therefore we sampled the size reduction parameter for each cell division from a Gaussian distribution of 160±20 nm (Fig. 1d, Fig. S3). The smallest viable diameter for *S. turris* is around 10 μm (Drebes, 1964; von Stosch and Drebes, 1964). Therefore, a cell that reached a diameter smaller than 10 μm was traded with a cell at the maximum size (54±1 μm), simulating size restoration via an auxospore. To mimic the experimental growth conditions the population divided synchronously, and after the seventh cell division a random pick of 10^6^ cells was done to restore the population back to its initial size, resembling the weekly culture dilution.

When running such a simulation, the mean cell size of the population shows a saw-tooth pattern caused by alternating periods of gradual reduction and more rapid size restoration, and the variance increases over time (Fig. 2 c). During the first 83 generations, the mean diameter decreases from 54±1 μm to 48.1±1.2 corresponding to an average reduction of 71 nm per generation. Note that the mean size reduction rate of the population (which is different from size reduction of each cell) is considerably slower than the experimental culture, indicating that the simulation does not capture every quantitative aspect of the culture behavior accurately.

Over 574 simulated generations, the mean diameter decreased to a minimum of 15.0±3.5 μm and the variance slightly increased. From that point, the population has a bimodal size distribution, marked by a broad variance. In the following generations, the mean diameter rapidly increases as with each cell division more cells reach a size of 10 μm, after which they recover to 54 μm. After each cycle, the mean cell size oscillates with a progressively smaller amplitude and eventually reaches a steady state that fluctuates around 32 μm, with a high variance (Fig. 2 d).

Using our experimental setup, we monitored the changes in cell size in six clonal cultures for a time period of more than a year (Fig. 3, Cultures 1-5 started independently and on day 60 of Culture 1 it was split into two replicates, 1a and 1b). Cultures 1a, 1b and 2, which were founded by large cells, with a diameter larger than 40 μm, initially decreased in size (Fig. 3, grey shaded areas), with a consistent slope of −135 to −125 nm per generation during this period, as determined by least-squares linear regression.

**Figure 3.**
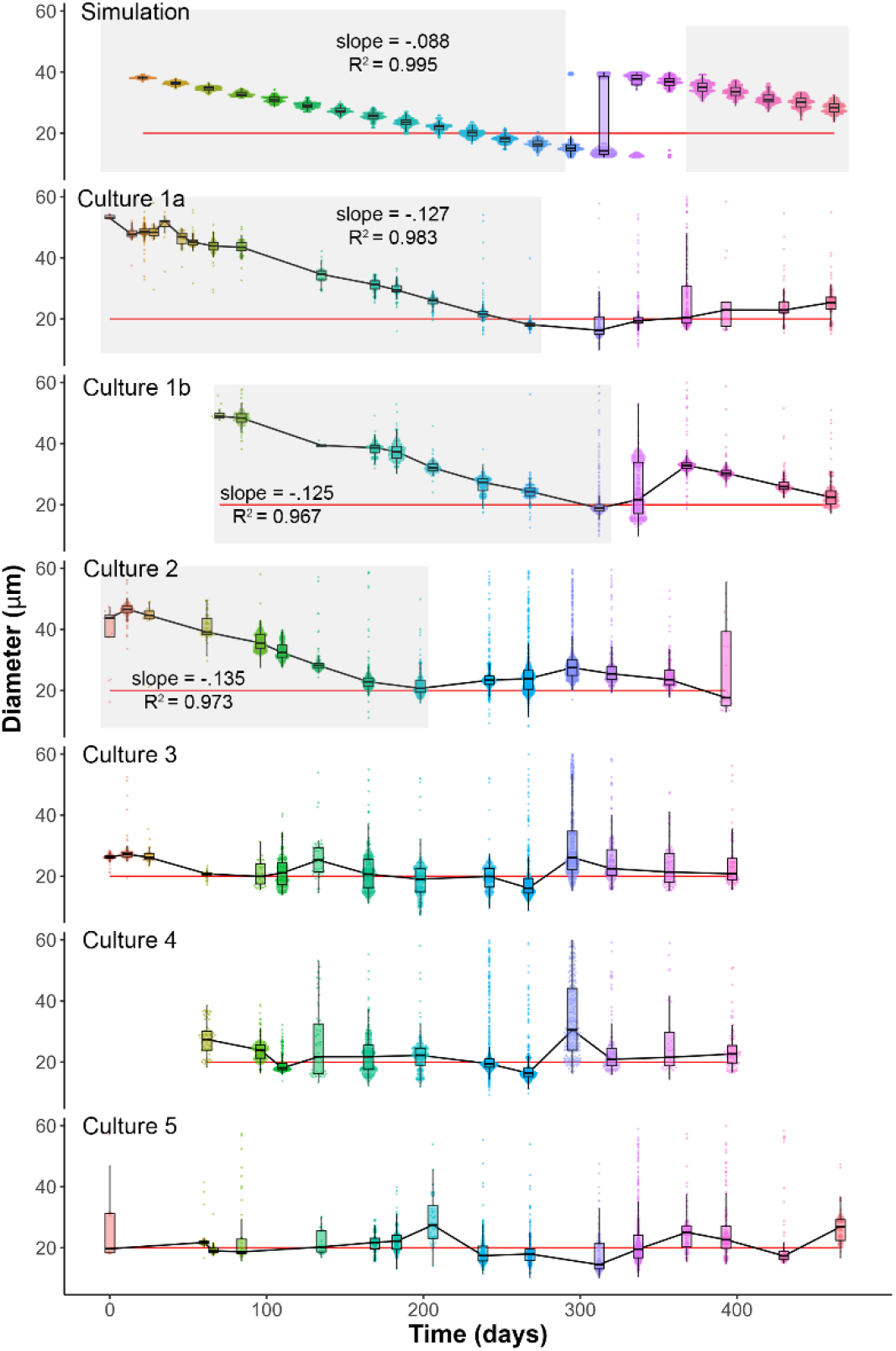
Change in size of *S. turris* cultures over long time periods. ‘Bee swarm’ plots and boxplots of cell diameter changes with generations in simulated dataset and six experimental cultures. Each point represents a single cell. Culture median cell diameter is shown as a line. Gray-shaded areas indicate periods of size reduction. For comparison with the experimental data, the simulation parameters were: Start with 10^6^ cells of 40±1 μm, size reduction per generation 200±20 nm.

Once the median cell size has decreased to about 20 μm (Fig. 3 red lines) the size reduction stalls in all cultures, which from then on show variable patterns of size changes. Notably, none of the large experimental cultures (1a, 1b and 2) recovered their size back to the original size of the founder cultures. Instead, the median cell size of most cultures fluctuates around 20 μm, with sizes increasing and decreasing to different extents and at different rates. Cultures 3 to 5 which were founded by narrow cells, with a diameter smaller than 25 μm fluctuated throughout the entire experiment. To test different environmental factors on cell size reduction, duplicates of the five clonal cultures 1 to 5 were grown at low light conditions. The cell division rate in low light was 10 times slower but the overall cell size change patterns were the same (Fig. S4).

Only one of the 6 cultures had a size change pattern that somehow resembles the simulated size restoration pattern (Fig. 3, Culture 1b). Culture 1b started with a median diameter of 48.9 μm and gradually decreased to 18.9 μm over 242 generations, corresponding to a reduction of 124 nm per generation. From there the unimodal distribution (mode 19 μm) changed to a bimodal distribution (modes 17 μm and 35 μm). At the next measurement the population of small cells had completely disappeared, resulting in a once again unimodal distribution with a median size of 32.8 μm (mode 31 μm). The variance of this culture also slightly resembles the expected pattern. However, even this culture did not restore the maximal cell size as predicted by the simulation and the other clonal cultures went through several small oscillations in median cell size while mainly remaining unimodal.

To test the robustness of the cell size change patterns that we found in the six long-term cultures, we isolated 20 additional clonal cultures and monitored their sizes weekly for a shorter time of 83 generations (Fig. 4, Fig. S5, Table S1). All of the 11 short-term cultures that were founded by large cells (median diameter ranging from 54.5 μm to 57.2 μm), showed a very robust gradual size reduction, slopes ranging from −155 to −128 nm per generation (Fig. 4a). On the contrary, cultures that were founded by smaller cells, with median diameters ranging from 17.7 μm to 36.2 μm, showed variable patterns (Fig. 4b). Of these cultures, three (S1, S3 and S4) showed a highly significant reduction in median size albeit with less steep slopes than the cultures in Fig. 4a. These three were among the 5 with the highest median start size. Culture S2 increased in size, with weak significance. The remaining four cultures did not significantly change in size. In general, these short-term cultures show similar behavior to the long-term experiments, where size reduction ceases once cells reach a size of 20 to 25 μm. From that point, instead of the expected sharp increase in cell size, most cultures develop a wider size distribution (see Fig. 3, Fig S5) with a median size that fluctuates between 20 and 25 μm.

**Figure 4.**
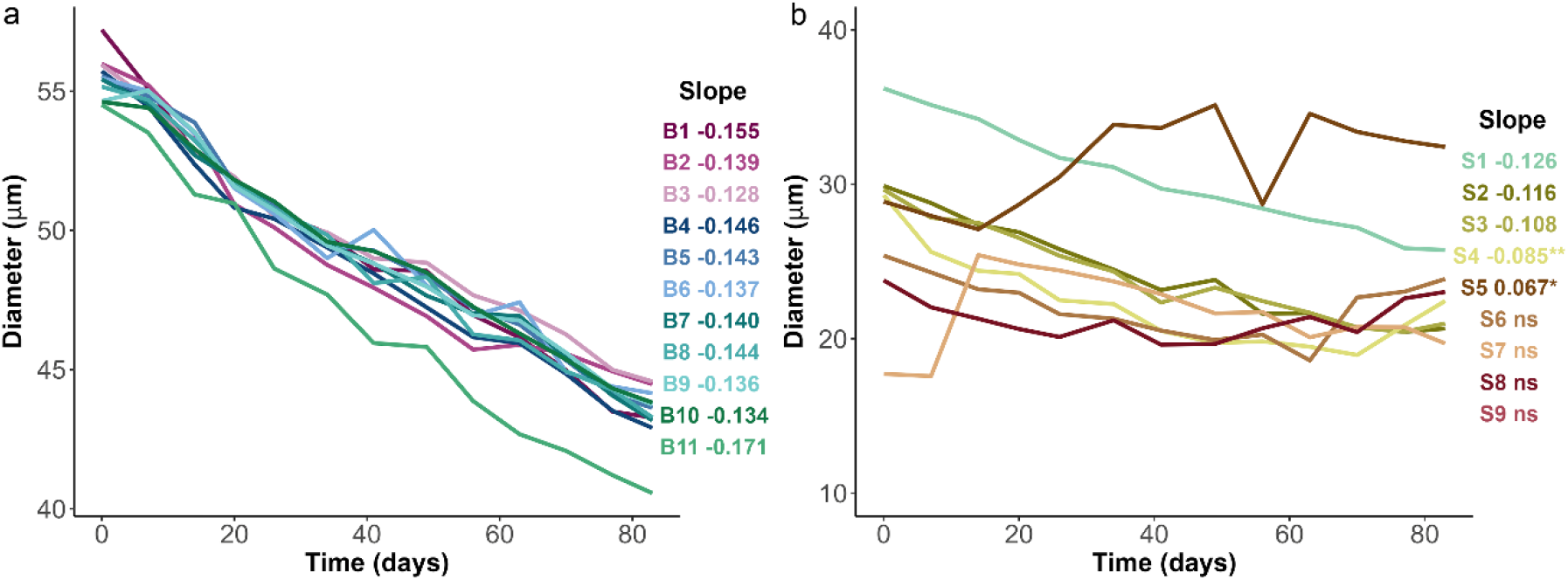
*S. turris* cell size patterns over short-term culturing. Progression of median cell diameter of *S. turris* cultures that started as homogeneous cultures with (**a**) median initial size of 53.9 μm and up, and (**b**) median initial size of 36.2 μm and lower. For each culture the slope of the least-squares linear regression line (i.e. size change per generation) is indicated, *p < 0.05, **p<0.01, and slopes without annotation p < 0.0001.

The size measurements suggest that smaller cells regulate their sizes via mechanisms that differ from the geometrical constraints of the MacDonald-Pfitzer model. We used visual inspection of *S. turris* cultures to monitor the observed patterns of cell size fluctuations. In the optical microscope it is impossible to measure the expected difference of ~150 nm between valves of the same cell. Nevertheless, occasionally we found cells with valves of outstandingly different sizes. In all such cases, the new valve was larger than the parental valve (Fig. 5a-b). Additionally, in some elongated cells that have not yet divided, the girdle band region is visibly wider than the valves (Fig. 5c-d). Thus, the girdle band region can be flexible to accommodate newly formed valves that are larger than the parental valves. On a finer scale, TEM images show parental valves tapering outward towards the valve rim, allowing the girdle band region and consequently the new valves to be considerably wider than the parental valve (Fig.5 e-f). Additionally, it appears that the attachment of the girdle bands to the valve sometimes leaves space for a new valve of the same size. Overall, even though the new valve can be smaller than the parental valve by twice the girdle band thickness, this is not a mandatory constraint that dictates cell size in *S. turris*.

**Figure 5.**
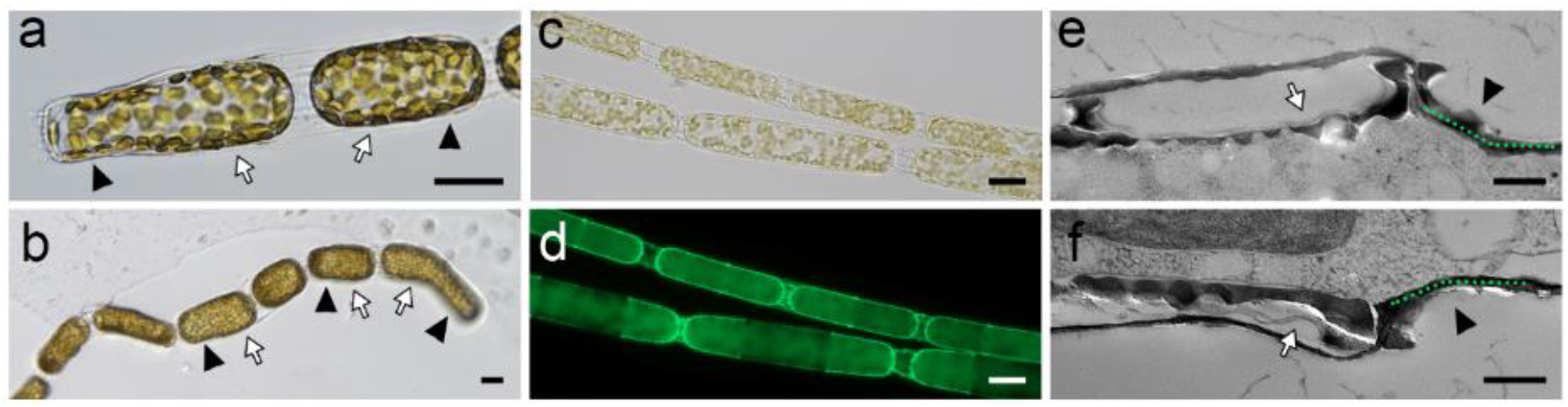
*S. turris* morphologies allowing cell size fluctuations. (**a-b**) Cells with larger new valve than parental valve. Scale bars: 20 μm. (**c-d**) Cells with girdle band region wider than the valves, imaged both in bright field (c), and with PDMPO fluorescence (d) that shows only the silica cell wall. Scale bars: 20 μm. (**e-f**) TEM micrographs of parental valves tapering out (green dashed lines) toward the rim. Black arrowheads: parental valve, white arrows: new valves. Scale bars: 500 nm.

## Discussion

The long-held paradigm of diatom cell size cycles, due to geometrical limitations imposed by the rigid silica cell walls, is backed by surprisingly little evidence for centric diatoms. For some highly silicified diatoms it is well documented that every vegetative cell division causes a decrease in cell size, and restoration to larger sizes requires a shift to sexual reproduction, but the situation for most diatoms, both in natural environments and in culture, is less clear. Our results from *S. turris* cultures, point to a more complex mechanism, in which geometrical size reduction plays a dominant role for large cells, but is overridden by other processes at smaller sizes. This is evident from both the highly reproducible rate of size decrease above a certain size threshold, as well as the small, highly variable, oscillations around a single value for small cells.

These results indicate a dual mechanism for cell size regulation, which is in agreement with numerous former cell size studies in centric diatoms. Our literature survey shows that natural diatom populations rarely show size distribution patterns that could arise purely from geometrical constraints (Table 1). The changes in cell size in a vast majority of the natural populations were associated with changes in environmental factors, demonstrating that cell size regulation in diatoms is flexible enough to allow adaptation to the environment. Accordingly, simulations that mimic the MacDonald-Pfitzer regime need to add some external inputs, along with imposing a limited lifespan for newly formed valves, in order to avoid the equilibrium state of an intermediate mean with a large variance, as seen also in our model, and reflect better the narrow size distributions found in natural and experimental diatom populations (Hense and Beckmann, 2015; Fuhrmann-Lieker *et al*., 2021).

Our well constrained experimental approach, based on controlled laboratory settings, can directly reveal size change patterns without the complications of environmental fluctuations or overlapping populations. There exist only a handful of examples of such studies for centric diatoms, and the predicted cycle of size decrease and recovery was reported in only two of them (Gross, 1937; Assmy *et al*., 2008). Interestingly, all other long-term controlled laboratory studies reported results similar to ours, i.e. conserved size reduction rates followed by periods of homeostasis with occasional jumps towards either larger or smaller cells (Findlay, 1969; Desikachary and Rao, 1973; Costello and Chisholm, 1981; Armbrust and Chisholm, 1992; Von Dassow, Chepurnov and Armbrust, 2006). Similar cell size homeostasis has also been reported for pennate diatoms, and notably, four of the most extensively studied species *Thalassiosira pseudonana, Phaeodactylum tricornutum*, *Cylindrotheca fusiformis* and *Navicula pelliculosa*, (Geitler, 1932; Wiedling, 1948; Chepurnov *et al*., 2008; Rose and Cox, 2013). Cell size homeostasis as observed in our laboratory cultures thus appears to be an important additional phase of the diatom life cycle.

Even though our high-throughput imaging approach monitored thousands of cells, we are not able to follow the size progression of each cell progeny. Occasionally, we observe the emergence of cells with sizes that are very different, and often much larger, from the ones in their mother culture. We hypothesize that there are several ways for this phenomenon to occur in our cultures. One of them is via auxospores, as cells with the structural characteristics of vegetative auxosporulation and sexual auxosporulation were sometimes seen. Though, we never observed an initial valve being formed within these putative auxospores. Another option is biological heterogeneity that leads to variable sizes in the culture. The sporadic appearance of large cells from a population of small cells is common, however, such events usually do not affect the population tendency to maintain within their optimal size range.

Cell size is a fundamental trait which directly affects physiological and ecological functions such as nutrient uptake, metabolic flux, buoyancy and susceptibility to predation (Marañón, 2015). Size plasticity is important in a changing environment as it can provide an adequate response to stresses such as nutrient limitation and high temperatures (Peter and Sommer, 2013; Hillebrand *et al*., 2022). However, under stable conditions unicellular organisms usually maintain a characteristic size, indicative of internal mechanisms that sense and regulate cell size (Jun *et al*., 2017; Schmoller, 2017). The diatom life cycle is uniquely characterized by long periods of progressive cell size reduction under stable environmental conditions. Since sexual reproduction usually produces offspring of the largest species-specific size and can only be induced above a specific size threshold it was proposed that the reduction-restitution cycle might act as an internal clock mechanism to accurately time and synchronize sex (Lewis, 1984). However, our investigations show that the size reduction period is commonly followed by a period of size homeostasis in which diatoms are able to maintain their cell size within a small range. Thus, timing of the sexual cycle and corresponding leap in cell size, appears to be predominantly triggered by environmental changes or encounters with potential mates, as is the case for other unicellular organisms (Moeys *et al*., 2016; Annunziata *et al*., 2022).

To conclude, these observations suggest that two distinct mechanisms regulate the diatom cell cycle, reduction and homeostasis. Large cells enter a controlled path of gradual size reduction towards a size range in which sexual reproduction can be induced by environmental changes. Once populations have reached a small size, the individual cells seem to employ a variety of mechanisms resulting in maintenance of an optimal size range.

## Acknowledgements

This project has received funding from the European Research Council (ERC) under the European Union’s Horizon 2020 research and innovation programme (grant agreement No. 848339).

## Data availability

The data supporting the findings of this study is available from the corresponding author upon reasonable request. The code used in this study is available through the following link: xxxx

