## Supporting information for "Decoupling cell size homeostasis in diatoms from the geometrical constraints of the silica cell-wall"

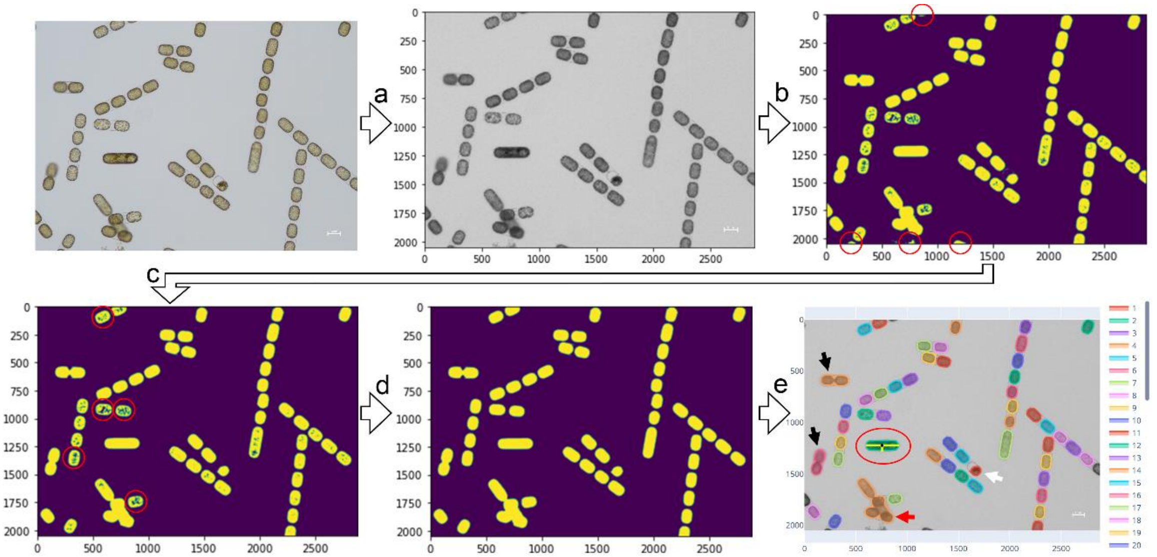


**Figure S1. Automated image processing. a)** RGB images are converted to grey-levels, **b)** and a threshold is applied to segment cells with lower pixel intensities from the background with higher pixel intensities. **c)** Regions below a certain size threshold are removed (encircled regions), **d)** and regions with holes below a certain size threshold are filled (encircled regions). **e)** Morphological properties of interest (e.g. minor and major axis length, yellow bars in encircled region) are measured for each the labeled regions. Some mislabeled regions were not filtered out: daughter cells that are not appropriately separated (black arrows), overlapping cells (red arrows) and damaged cells/impurities (white arrow). A majority of such mislabeled regions are later removed from the data by applying an upper (60 µm) and lower (8 µm) threshold for the minor axis length.

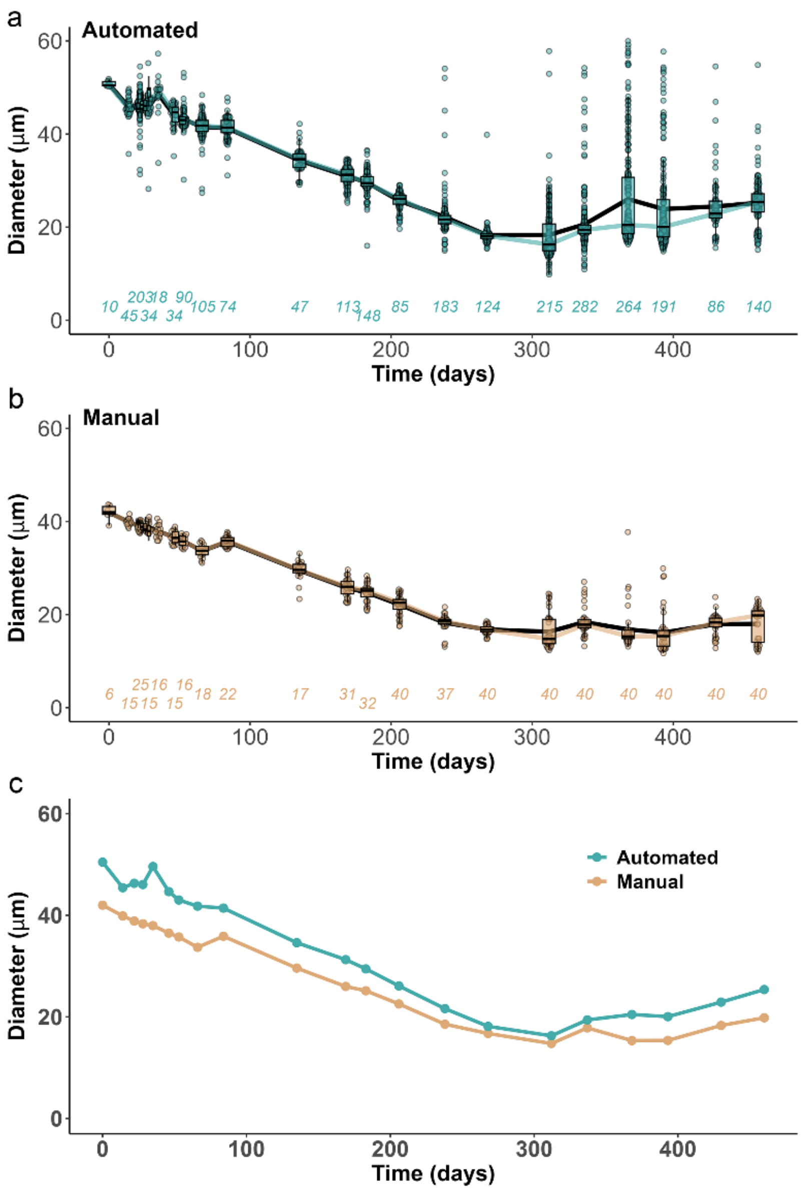


**Figure S2. Change in size of an *S. turris* culture measured using automated and manual approach. a)** 'Bee swarm' plots and boxplots of cell diameter per generation measured using automation, and **b)** manually. Each point represents a single cell. Colored and black lines connect the median and mean diameter measured at each time point respectively. Sample size for each measurement is indicated at the bottom of the plot in italics. **c)** Median cell diameter over time.

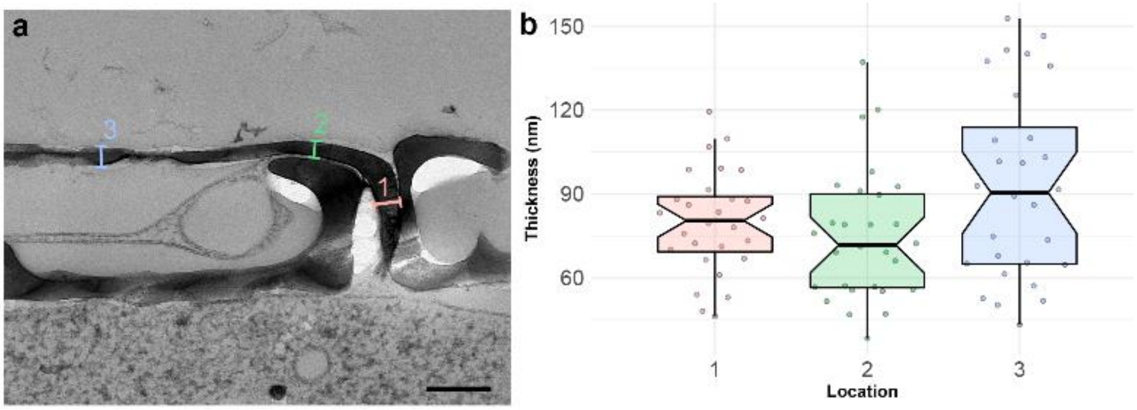


**Figure S3. Thickness of *S. turris* girdle bands.** (**a**) TEM micrograph of the valve-valve boundary, indicating measurement locations. Scale bar: 200 nm (**b**) Thickness of 28 girdle bands, measured at 3 locations.

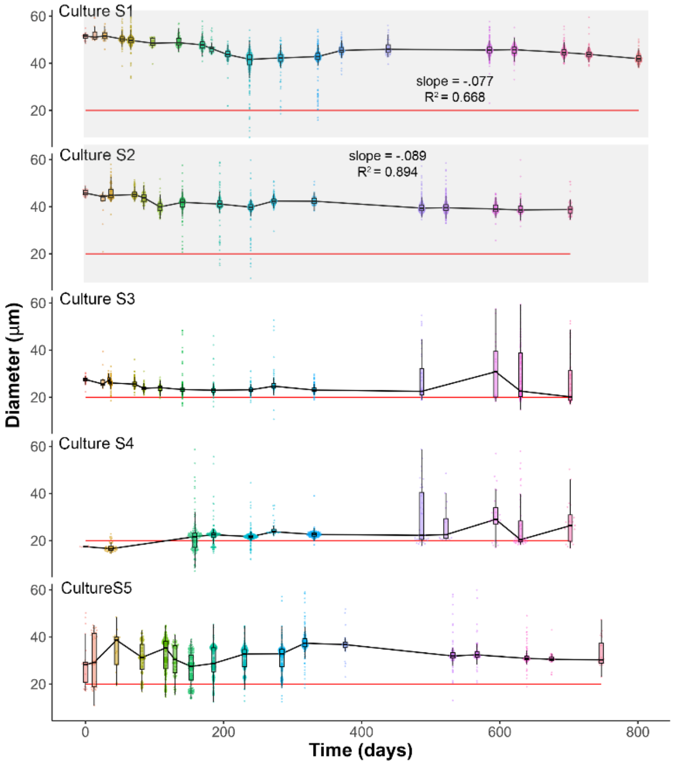


**Figure S4. Change in size of *S. turris* cultures over long time periods at low light.** ‘Bee swarm’ plots of cell diameter by generation in five experimental cultures. Each point represents a single cell. Culture mean cell diameter ±1 standard deviation are shown as a line and shaded area. Gray-shaded areas indicate periods of size reduction with continuous size reduction.

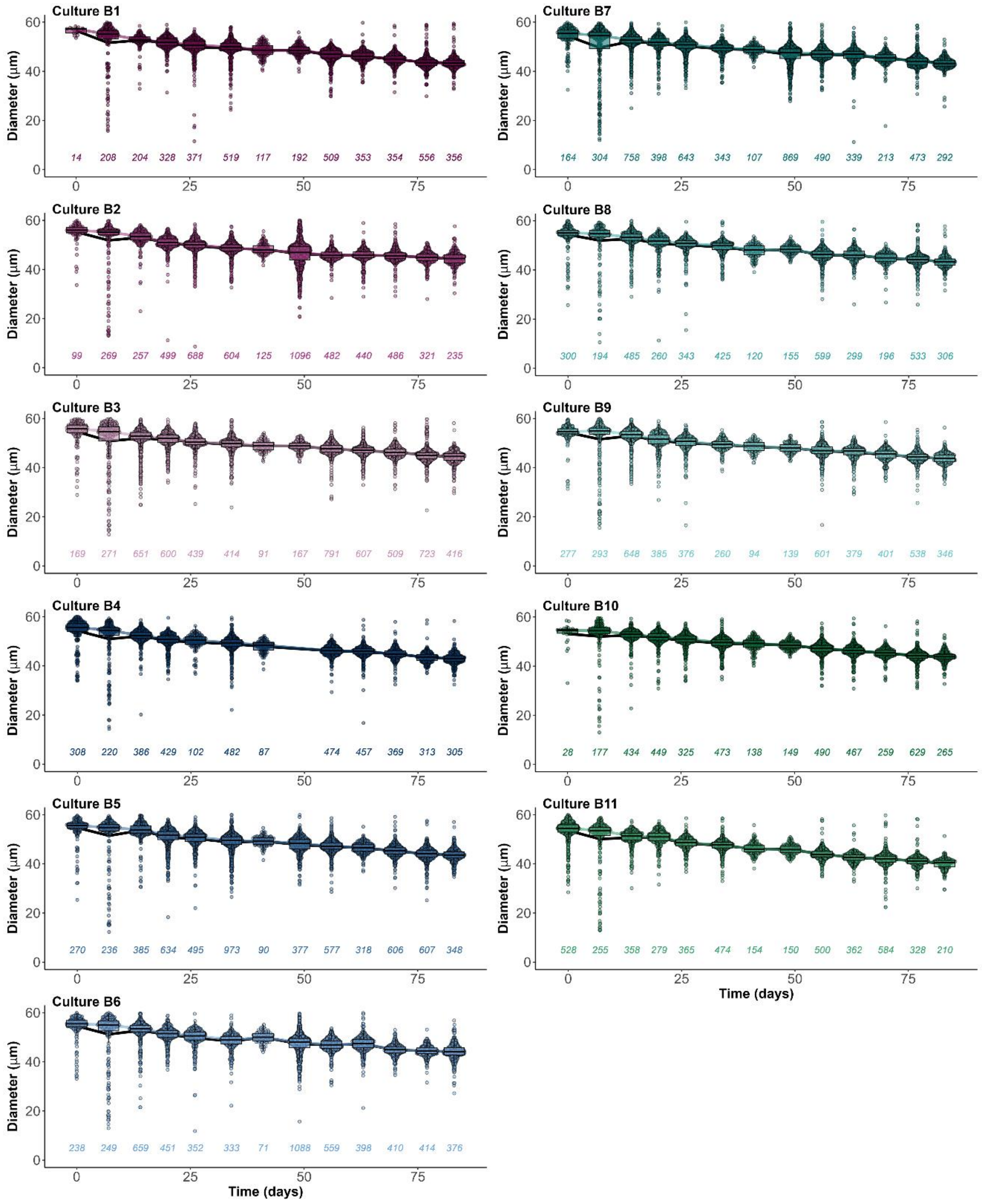


**Figure S5. Change in size of *S. turris* cells over short-term culturing.** 'Bee swarm' plots and boxplots of cell diameter change with time in 20 experimental cultures. Each point represents a single cell. Culture median and mean cell diameter are shown as colored and black lines, respectively. Numbers below the plot indicate sample size for each time point.

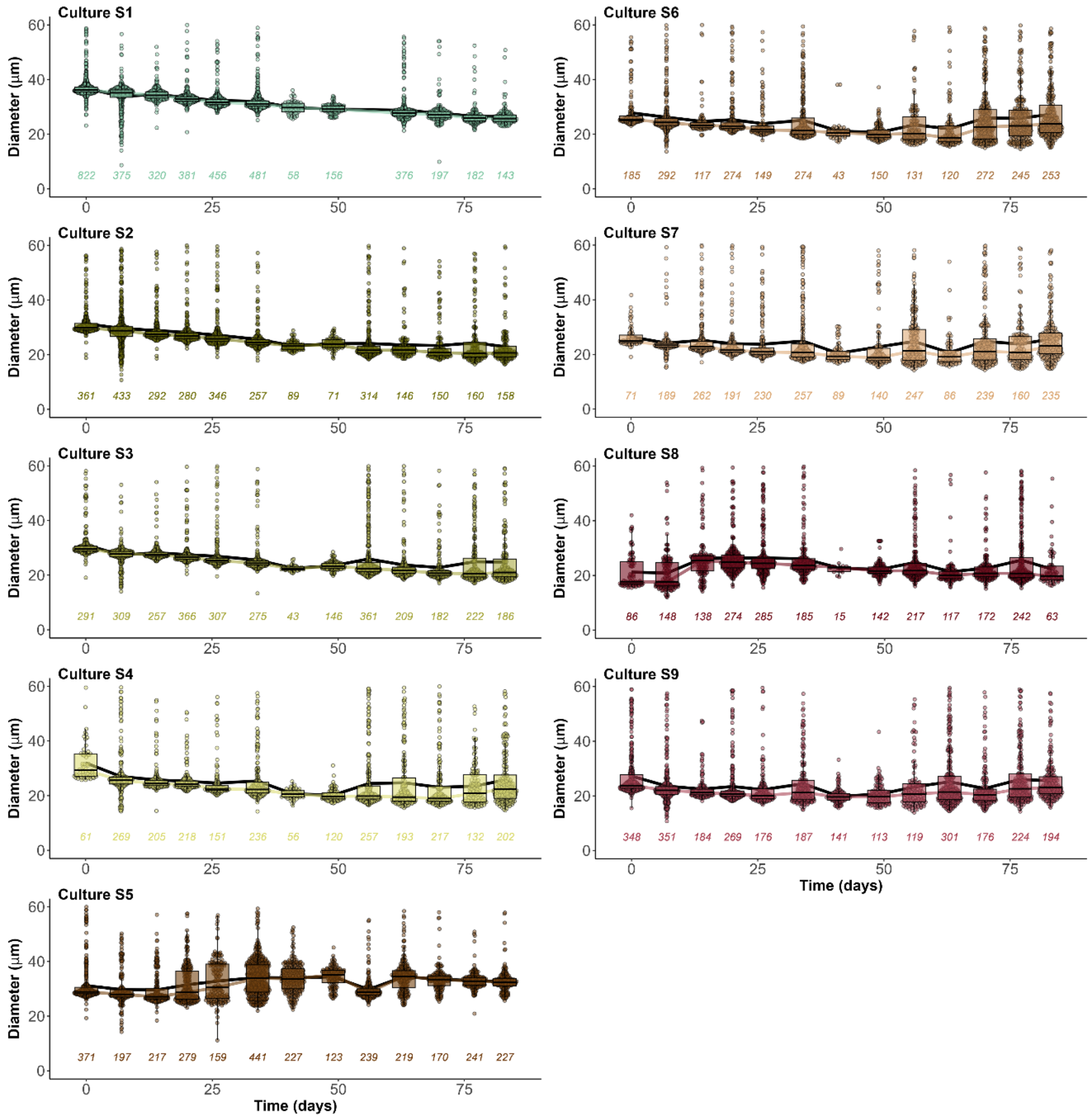


**Figure S5 - continued**

### Table S1 – Cell size change summary statistic of *S. turris* cultures.

| Culture | $R^2$ | Adj. $R^2$ | F | df | Res. Std | Intercept (µm) | Slope (µm/gen.) | p |
|---|---|---|---|---|---|---|---|---|
| **Culture B1** | **0.9728** | **0.9704** | **394** | **1, 11** | **0.7396** | **55.63** | **-0.155** | **<0.000** |
| **Culture B2** | **0.9237** | **0.9167** | **133.1** | **1, 11** | **1.138** | **54.68** | **-0.139** | **<0.000** |
| **Culture B3** | **0.9729** | **0.9704** | **394.4** | **1, 11** | **0.6112** | **54.89** | **-0.128** | **<0.000** |
| **Culture B4** | **0.9811** | **0.9793** | **520.2** | **1, 11** | **0.6032** | **54.72** | **-0.146** | **<0.000** |
| **Culture B5** | **0.9851** | **0.9838** | **729.2** | **1, 11** | **0.5013** | **55.23** | **-0.143** | **<0.000** |
| **Culture B6** | **0.9623** | **0.9589** | **281** | **1, 11** | **0.78** | **55.01** | **-0.137** | **<0.000** |
| **Culture B7** | **0.9857** | **0.9844** | **759.4** | **1, 11** | **0.481** | **54.89** | **-0.140** | **<0.000** |
| **Culture B8** | **0.9852** | **0.9839** | **733.6** | **1, 11** | **0.5028** | **54.95** | **-0.144** | **<0.000** |
| **Culture B9** | **0.9789** | **0.977** | **511** | **1, 11** | **0.5698** | **54.82** | **-0.136** | **<0.000** |
| **Culture B10** | **0.9939** | **0.9933** | **1791** | **1, 11** | **0.2985** | **54.71** | **-0.134** | **<0.000** |
| **Culture B11** | **0.9832** | **0.9816** | **641.9** | **1, 11** | **0.6378** | **53.92** | **-0.171** | **<0.000** |
| **Culture S1** | **0.9838** | **0.9822** | **607.8** | **1, 11** | **0.4773** | **35.64** | **-0.126** | **<0.000** |
| **Culture S2** | **0.956** | **0.952** | **239.2** | **1, 11** | **0.7111** | **29.08** | **-0.116** | **<0.000** |
| **Culture S3** | **0.9358** | **0.9299** | **160.2** | **1, 11** | **0.8041** | **28.56** | **-0.108** | **<0.000** |
| **Culture S4** | **0.6152** | **0.5803** | **17.59** | **1, 11** | **1.918** | **25.85** | **-0.085** | **0.0015** |
| **Culture S5** | **0.4282** | **0.3762** | **8.237** | **1, 11** | **2.213** | **28.57** | **0.067** | **0.01524** |
| Culture S6 | 0.1425 | 0.0645 | 1.827 | 1, 11 | 1.887 | 23.25 | -0.027 | 0.2036 |
| Culture S7 | 0.2035 | 0.131 | 2.81 | 1, 11 | 1.674 | 22.59 | -0.030 | 0.1219 |
| Culture S8 | 0.01366 | -0.076 | 0.1524 | 1, 11 | 2.636 | 22.09 | -0.011 | 0.7037 |
| Culture S9 | 0.00202 | -0.0887 | 0.02221 | 1, 11 | 1.345 | 21.36 | -0.002 | 0.8842 |
